# In situ genetically targeted chemical assembly of polymers on living neuronal membranes

**DOI:** 10.1101/2022.12.27.521974

**Authors:** Anqi Zhang, Kang Yong Loh, Chandan S. Kadur, Lukas Michalek, Jiayi Dou, Charu Ramakrishnan, Zhenan Bao, Karl Deisseroth

## Abstract

Multicellular biological systems, most notably living neural networks, exhibit highly complex physical organization properties that pose challenges for building cell-specific and biocompatible interfaces. We developed a novel approach to genetically program cells to chemically assemble artificial structures that modify the electrical properties of neurons *in situ*, opening up the possibility of minimally-invasive cell-specific interfaces with neural circuits in living animals. However, the efficiency and biocompatibility of this approach were challenged by limited membrane targeting of the constructed material. Here, we report a method with significantly improved molecular construct properties, which expresses highly localized enzymes targeted to the plasma membrane of primary neurons with minimal intracellular retention. Polymers synthesized *in situ* by this approach form dense clusters on the targeted cell membrane, and neurons remain viable after polymerization. This platform can be readily extended to incorporate a broad range of materials onto the surface membranes of specific cells within complex tissues, using chemistry that may further enable the next generation of interfaces with living biological systems.

## Main

Multicellular biological systems can display highly complex structural and organizational properties, posing challenges for investigators seeking to achieve cell-specific electrical interfaces without inducing damage. For example, the human brain contains nearly 100 billion neurons and over 100 trillion synaptic connections, and the basic elements crucial to neuronal communication and signal propagation (e.g., synapses and ion channels) are on the nanometer scale. Despite decades of efforts in shrinking bioelectronic devices to the nanometer scale^1^ and scaling up the fabrication process, current devices designed for the brain can only interface with up to hundreds of cells at a time^2^, and are unable to achieve seamless integration with the biological systems with cell-type specificity. A new approach to tackle this problem is by genetically programing specific cells within intact biological systems to build artificial structures with desired form and function *in situ*. It had previously been shown that conductive polymers can be directly synthesized on cells in living tissue with electrochemical polymerization^3^, or in organisms and tissues with native oxidative environments^4–6^ or oxidative enzymes^7^. However, none of these approaches enabled the critical goal of cellular specificity. We have taken the first step in this direction with the launch of the new field of genetically targeted chemical assembly (GTCA)^8^, which uses cell-specific genetic information to guide neurons to initiate deposition of polymer materials *in situ* with a variety of electrical conduction properties. Specifically, an ascorbate peroxidase Apex2^9^ was expressed in neurons as the catalyst, and was used to initiate hydrogen peroxide (H_2_O_2_)-enabled oxidative polymerization of either conductive polymers and insulating polymers. Electrophysiological and behavioral analyses were used to confirm that the genetically targeted assembly of functional polymers remodeled membrane properties and modulated cell type-specific behaviors in freely moving animals.

Despite this initial success, the procedures previously used for the proof-of-concept system displayed a major limitation: the Apex2 was not specifically targeted to the plasma membrane alone; the majority of Apex2 was retained intracellularly (Supplementary Fig. S1). Developing a process that could efficiently place the reaction centers fully in the extracellular space, on the external side of the membrane, will be critical to further applications of this biomanufacturing platform for the following reasons. First, live cells with intact membrane will not typically be permeable to polymer precursors or materials, so insufficient membrane-display of enzymes may lead to a low yield for the chemical synthesis in living systems. Second, increasing the number of enzymes catalyzing the reactions (leveraging extracellular space) may allow reduced concentrations of reaction-condition components (such as H_2_O_2_) and thereby improve biocompatibility. Third, localizing reactions to extracellular space may limit adverse effects on native intracellular chemistry. Indeed, previous reports have demonstrated that certain intracellular polymerization reactions may be toxic to cells and can induce apoptosis^5,10,11^. Finally, beyond membrane localization, many other avenues for advancing the technology also exist; for example, Apex2 is not optimized for applications on the cell surface. Another well-characterized peroxidase, horseradish peroxidase (HRP), catalyzes the same reactions as Apex2 with faster kinetics and greater resistance to H_2_O_2_-induced inactivation^12^.

Here, we present a next-generation GTCA for targeted polymer assembly with highly localized HRPs on the plasma membrane of primary neurons with minimal intracellular retention. Polymers synthesized by this approach form dense clusters around the living neuron membrane of interest, and all neurons remained viable after polymerization.

### *In situ* genetically targeted chemical assembly of polymers on the surface of neurons

The schematic in Fig. 1a provides an overview of our next-generation method. We first designed a template plasmid DNA construct for expressing membrane-displayed HRP in primary neurons. The new construct encodes the neuron-specific human Synapsin (hSyn) promoter, followed by an IgK leader sequence that initially directs the protein into the endoplasmic reticulum (ER), FLAG tags for antibody detection, HRP, a transmembrane (TM) domain as the membrane targeting anchor, 2A self-cleaving peptides, and enhanced yellow fluorescent protein (YFP). The targeted neurons thus are anticipated to express membrane-displayed HRP and cytosolic YFP. HRP location can be determined by staining with antibodies targeting the FLAG tags. Upon addition of small-molecule polymer precursors and H_2_O_2_, the membrane-displayed HRPs act as reaction centers, facilitating oxidative radical polymerization on targeted neurons. Because of the low solubility of the resulting polymers, these synthesized polymers are expected to be deposited onto the targeted cell membrane. Here, we selected two polymer precursors, N-phenyl-p-phenylenediamine (aniline dimer) and 3,3’-diaminobenzidine (DAB monomer), for the deposition of a conductive polymer, polyaniline (PANI), and a non-conductive polymer, poly(3,3’-diaminobenzidine) (PDAB), respectively (Fig. 1b). Both reactions occur in biocompatible aqueous solutions with 0.05 mM H_2_O_2_.

**Figure 1.**
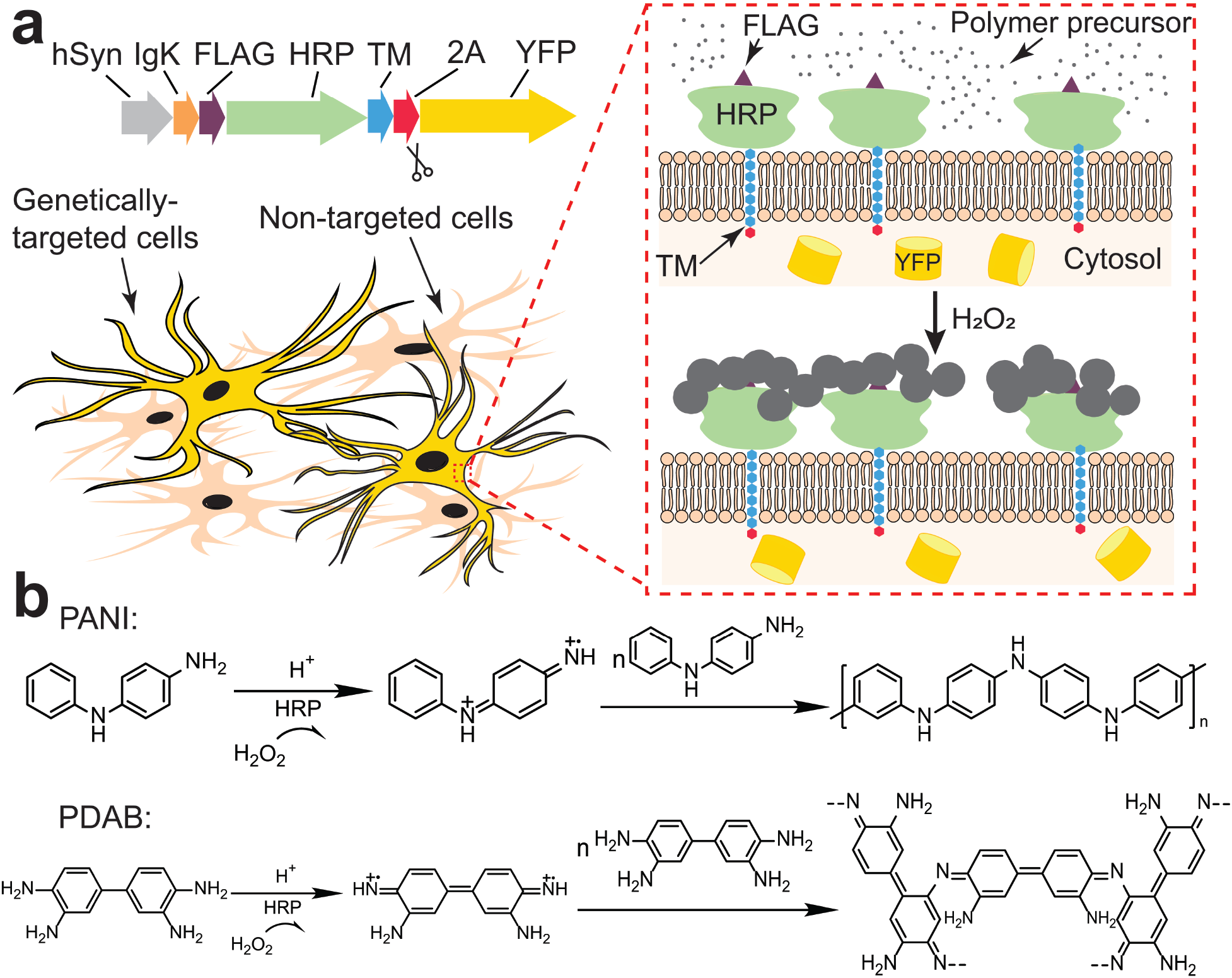
*In situ* genetically targeted chemical assembly of polymers on the surface of neurons. (**a**) Top, DNA construct backbone for expressing membrane-displayed HRP. The construct contains a neuron-specific hSyn promoter, followed by sequences coding for the IgK leader sequence, FLAG tags for antibody detection, HRP, a transmembrane (TM) domain as the membrane targeting anchor, 2A self-cleaving peptides, and enhanced yellow fluorescent protein (YFP). The targeted cells are expected to express membrane-displayed HRP and cytosolic YFP. Bottom, polymer assembly occurs specifically on the membrane of enzyme-targeted cells expressing cytosolic YFP. Inset shows HRP anchored on the cell membrane enabling extracellular HRP/H_2_O_2_-catalyzed polymerization. Polymer precursors form dark-colored aggregates deposited on the cell surface. Cells in beige are non-enzyme-targeted cells. (**b**) HRP-mediated oxidative polymerization of PANI (top) and PDAB (bottom) from polymer precursors: N-phenyl-p-phenylenediamine (aniline dimer) and 3,3’-Diaminobenzidine (DAB), respectively.

### Membrane localization of HRP and evaluation of peroxidase activity

Native transmembrane proteins are synthesized in the ER, transported to the Golgi apparatus, and eventually reach the plasma membrane. Building upon the construct backbone described in Fig. 1, we sought to test a panel of native transmembrane domains as membrane targeting anchors, to most efficiently recruit the native membrane trafficking machinery in primary neurons. We tested the transmembrane domains of two T-cell surface glycoproteins, CD8α^13^ and CD2^14^, for expressing both HRP and Apex2 (Methods); to quantitatively compare membrane expression profiles of these constructs, we performed antibody staining procedures involving non-detergent-permeabilized or detergent-permeabilized cells (Fig. 2a; Methods). Specifically, HRP or Apex2 fused to FLAG tags are detected with primary antibodies targeting FLAG, which then can be labeled with secondary antibodies conjugated with Alexa 647 fluorophores. Under these conditions, in non-detergent-permeabilized cells chiefly membrane-anchored enzymes can be optimally detected, whereas in detergent-permeabilized cells both intracellular and extracellular enzymes are robustly labeled.

**Figure 2.**
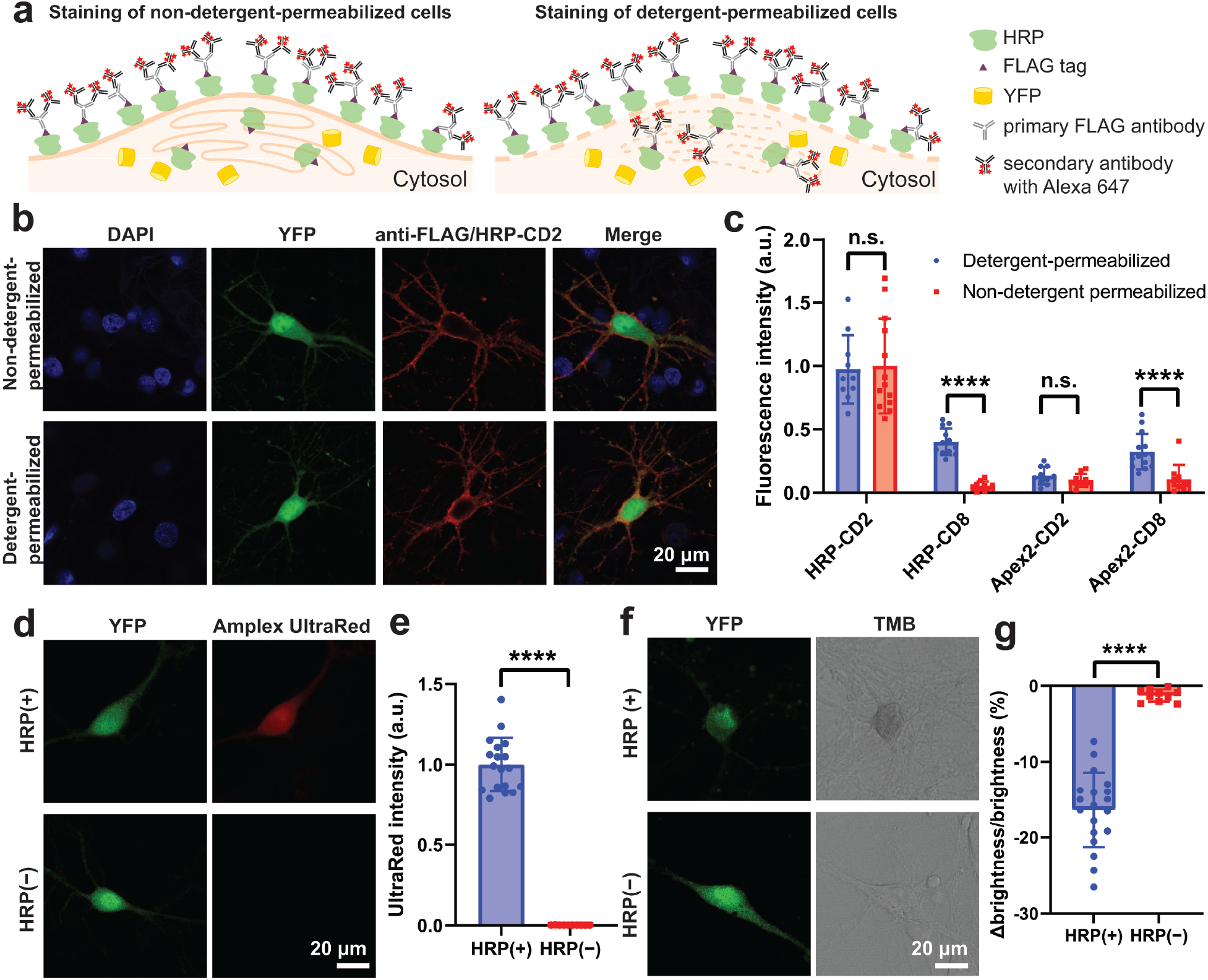
Membrane localization of peroxidases and evaluation of peroxidase activity. (**a**) Schematic: quantitative evaluation of membrane expression in non-detergent-permeabilized and detergent-permeabilized cells. HRP enzymes fused to the FLAG tag are detected with primary antibodies targeting FLAG, which are then labeled with secondary antibodies with Alexa 647 fluorophores. In non-detergent-permeabilized cells, only membrane-anchored HRP can be detected. In detergent-permeabilized cells, both intracellular and extracellular HRP are labeled. (**b**) Confocal microscopy of expression and membrane localization of HRP-CD2 in neurons. Four days post-transfection, cells were fixed and either permeabilized or not; HRP localization in HRP(+) cells (here, cells expressing cytosolic YFP) was detected by immunostaining. Cell nuclei were labeled with DAPI. The similarity in HRP localization in detergent-permeabilized and non-detergent-permeabilized cells, along with the uniform membrane-associated antibody fluorescence, demonstrate robust membrane-trafficking enabled by the CD2 membrane anchor. (**c**) Fluorescence intensity comparison of antibody-stained detergent-permeabilized vs. non-detergent-permeabilized neurons, expressing HRP or Apex2, with membrane anchors CD2 or CD8 (N = 10 cells each). Laser intensity and microscope settings were maintained consistent across conditions. (**d**) Evaluation of oxidative activity of HRP in living neurons expressing HRP-CD2 with the fluorogenic membrane-permeable dye Amplex UltraRed. The red fluorescent product of Amplex UltraRed oxidation indicates peroxidase activity. (**e**) Statistical comparison of the UltraRed intensity; N = 18 cells for HRP(+), N = 10 cells for HRP(−). (**f**), Evaluation of oxidative activity of HRP in fixed neurons expressing HRP-CD2 with the chromogenic substrate TMB; the blue reaction product (appearing dark in bright field images) indicates peroxidase activity. (**g**) Statistical comparison of the ratio (expressed as %) of brightness difference between neuron and background (“Δbrightness”), compared to background brightness. N = 19 cells for HRP(+), N = 10 cells for HRP(−). Values plotted are means ± s.d.; n.s. = nonsignificant, **** = P < 0.0001; two-tailed, unpaired, t-test.

Four days post-transfection with plasmid DNA, enzyme localization in HRP(+) and Apex2(+) cells (i.e. here, cells expressing cytosolic YFP) was detected by immunostaining, and cell nuclei were labeled with 4’,6-diamidino-2-phenylindole (DAPI). Representative confocal microscopy imaging of the HRP-CD2 neurons (Fig. 2b) revealed uniform membrane-associated antibody fluorescence on the soma and neurites in the anti-FLAG/Alexa 647 imaging channel. Similarity in HRP localization in detergent-permeabilized and non-detergent-permeabilized cells confirmed good membrane-trafficking properties of CD2 under these conditions. We compared the fluorescence intensity of antibody staining in neurons expressing HRP-CD2, HRP-CD8, Apex2-CD2, and Apex2-CD8 (N = 10 cells each; Fig. 2c); the percentage of enzyme-targeted immunofluorescence that was membrane-displayed (relative to total expression levels) was approximately 100%, 13%, 72%, and 25%, respectively. CD2 thus exhibited more efficient membrane-anchoring properties than CD8, and furthermore the overall expression level of HRP-CD2 was higher than that of Apex2-CD2. Indeed, polymerization results confirmed that reactions on live HRP-CD2 neurons were much faster than on Apex2-CD2 neurons (Supplementary Fig. S2). We therefore used HRP-CD2 neurons as HRP(+) cells for the rest of the present work, with YFP-CD2 neurons as negative control HRP(−) cells.

Next, we tested HRP activity in live and fixed neurons using two peroxidase substrates: Amplex UltraRed and 3,3’,5,5’-Tetramethylbenzidine (TMB), respectively. Amplex UltraRed is a fluorogenic substrate for HRP that produces bright red membrane-permeable product after oxidation^15^; fluorescence images confirmed that HRP(+) but not HRP(−) neurons turned red after reaction (Fig. 2d,e). TMB is a chromogenic substrate, which forms a water-soluble blue reaction product upon oxidation; bright field images confirmed that the coloration change only occurred in HRP(+) neurons (Fig. 2f,g).

### Morphological and mechanical characterizations of *in situ* deposited polymers

Next, we performed polymerization reactions of PANI and PDAB on HRP(+) and HRP(−) neurons (see Methods for reaction conditions). Bright field images showed that HRP(+) neurons exhibited dark-colored reaction products, but HRP(−) neurons did not (Fig. 3a). Reaction progress was quantified as the decrease in brightness of cells relative to background (Fig. 3b). PANI formed densely-distributed aggregates, while PDAB formed a thin uniform coating with scattered non-uniform aggregates. Scanning electron microscopy (SEM) imaging confirmed these morphologies of both polymer depositions (Fig. 3c, Supplementary Fig. S3, S4). HRP(+)/PANI neurons formed dense clusters around the neuronal soma and neurites, with average particle size of 120 ± 13 nm. By contrast, HRP(+) cells not exposed to PANI exhibited smooth membrane surfaces. The HRP(+)/PDAB neurons formed a thin layer with non-uniform aggregates of clusters, with average particle size of 115 ± 11 nm (Supplementary Fig. S4). Further comparison with the SEM images of HRP(−) cells confirmed localization specificity of this genetic targeting approach (Supplementary Fig. S4). We also used atomic force microscopy (AFM) to characterize the height and Derjaguin-Muller-Toporov (DMT) modulus mappings of PANI particles on the neuronal membrane (Fig. 3d, see Supplementary Fig. S5 for original images). Height maps of HRP(+)/PANI cells again showed clear distribution of ~100 nm polymer particles, similar to those seen in SEM images and uniformly deposited across the membrane. A significant increase was observed in the modulus maps after polymerization, with polymer particles showing modulus values at least three orders of magnitude higher than on cells without polymerization (Supplementary Fig. S5). Notably, the membrane regions between particles also showed much higher modulus than the pristine membrane, indicating that there might be a thin layer of PANI uniformly coated on cells under the particles. We plan to perform additional AFM measurements in the future to further characterize this polymer layer, and study how the membrane modulus increase changes the cell properties.

**Figure 3.**
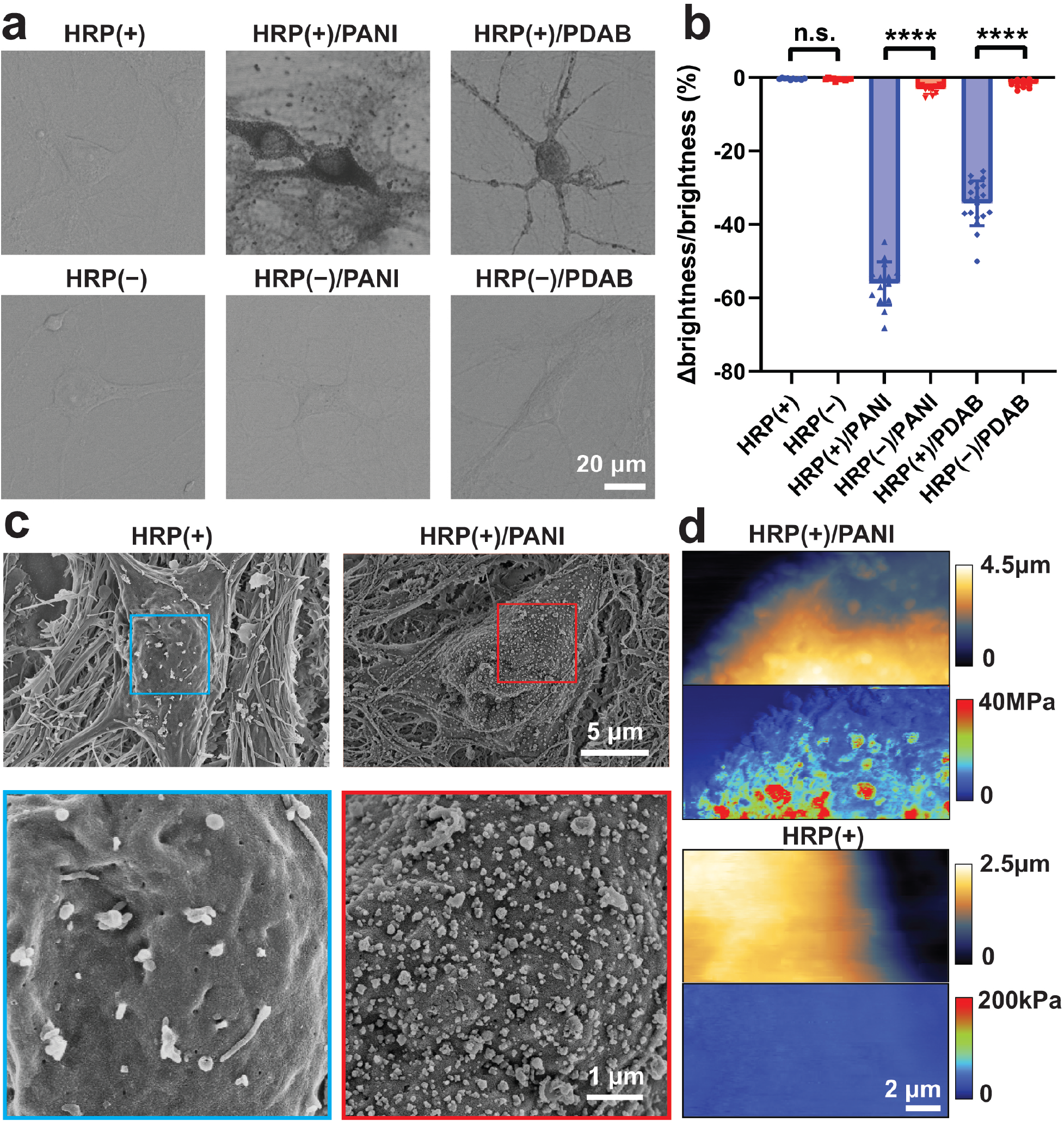
Morphological and mechanical characterizations of *in situ* deposited polymers. (**a**) Bright field images of live neurons with and without polymer deposition. (**b**) Statistical comparison of the ratio (expressed as %) of brightness difference between neuron and background (“Δbrightness”), compared to background brightness. N = 10 cells for HRP(+), N = 11 cells for HRP(−), N =15 cells for HRP(+)/PANI, N =11 cells for HRP(−)/PANI, N =19 cells for HRP(+)/PDAB, N = 13 cells for HRP(−)/PDAB. Values shown are means ± s.d.; n.s. = non-significant, **** = P < 0.0001; two-tailed, unpaired, t-test. (**c**) Representative SEM images of neurons with and without polymer deposition. HRP(+)/PANI show dense polymer aggregates on the membrane. (**d**) Representative AFM images of HRP(+) neurons with and without PANI deposition. Top, height channel; bottom, modulus map. The polymer particles exhibited significantly higher modulus compared to the membrane.

### Spectroscopic characterizations of *in situ* deposited polymers

To characterize the chemical composition of *in situ* deposited polymers, we used ultraviolet-visible (UV-Vis) absorption spectroscopy to compare with spectra previously reported for PANI and PDAB (Figs. 4a,b). HRP(+)/PANI spectra exhibit an absorption peak at approximately 370 nm that is attributable to π–π* transition of the benzenoid ring. The peak at 536 nm corresponds to intramolecular transition of benzenoid rings into quinoid rings^16,17^. The HRP(+)/PDAB spectra also exhibits an absorption peak at 370 nm, and the 460 nm peak corresponds to the characteristic absorption of PDAB^18^. Finally, confocal Raman microscopy was used to examine single cells with and without PANI (Fig. 4c) and PDAB (Supplementary Fig. S6). The bands at 830, 1000 and 1030 cm^−1^ correspond to the in-plane deformation of the rings. The bands at 1230, 1260, 1343, and 1444 cm^−1^ are attributable to the C–N and C=N stretching vibrations. The 1180 and 1510 cm^−1^ are from deformation vibrations of C–H and N–H, and the 934 and 1610 cm^−1^ correspond to the stretching vibration of C–C^19^. Together, these results confirmed that PANI was synthesized at the plasma membrane surface (Fig. 4c).

**Figure 4.**
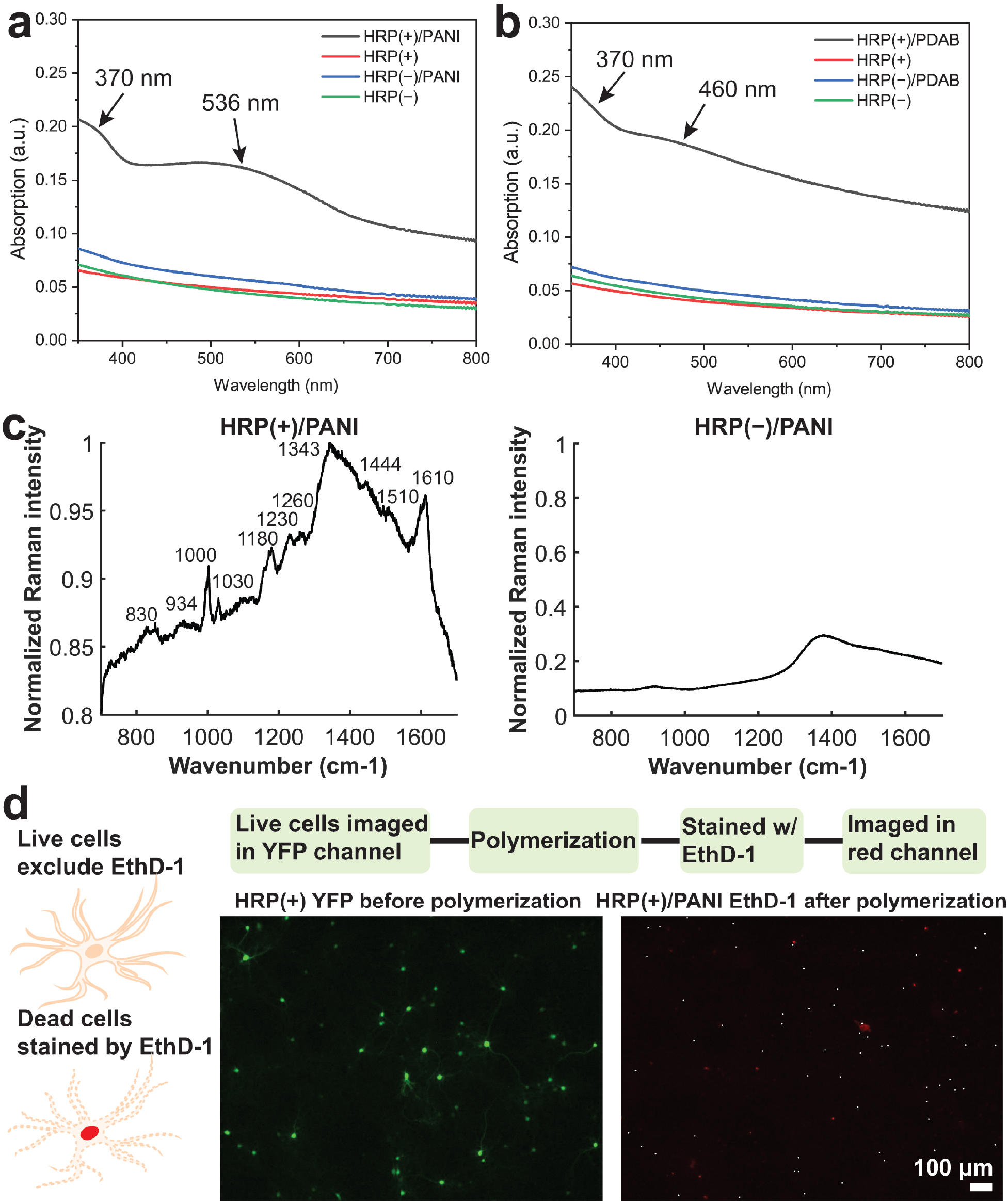
Spectroscopic characterizations of *in situ* deposited polymers and cell viability assay. (**a, b**) Normalized UV-Vis spectra of HRP(+) and HRP(−) neurons with and without PANI and PDAB deposition. Arrows indicate absorption peaks. (**c**) Raman spectroscopy analysis of the HRP(+) and HRP(−) after PANI deposition. (**d**) Cell viability test of polymerization conditions. Live HRP(+) cells were imaged with YFP, and after PANI deposition, were stained with ethidium homodimer (EthD-1), a red fluorophore that selectively stains dead cells with damaged membranes. White dots in the EthD-1 image mark locations of the HRP(+) cells. No white dot overlays with the red cells, indicating HRP(+) neurons remain viable after polymerization.

### Cell viability determination

We used a cell viability assay to monitor HRP(+) cells before and after polymerization (Fig. 4d, Supplementary Fig. S7, S8; see Methods). The YFP signals of live HRP(+) cells were obtained, and after PANI and PDAB deposition cells were stained with ethidium homodimer (EthD-1, a red fluorophore that only stains nucleic acids in dead cells). The locations of HRP(+) cells are marked in the EthD-1 channel with white dots (Fig. 4d); no white dot overlays with red cells, indicating all HRP(+) neurons remain viable after polymerization. In comparison, the positive control for EthD-1 staining on fixed dead cells confirmed that all dead cells were properly labeled using the same procedure (Supplementary Fig. S7); this cell viability test was repeated on N = 5 coverslips for each polymerization condition (Supplementary Fig. S8).

### Conclusion and outlook

Here we have addressed the key limitation of a new GTCA platform using genetically-targeted cells within living systems to instruct synthesis of functional materials, by deploying a method that allows precise targeting of enzymes onto the plasma membrane of primary neurons. Polymers synthesized *in situ* on neurons formed dense clusters on the membrane, and neurons retained viability post-reaction.

We anticipate that our approach will show generalized utility across a variety of application domains. First, targeted deposition of conductive or insulating polymers on living neuronal membranes, which increase or decrease capacitance respectively, will modulate excitability^8^ and translate into altered behavioral responses in a living animal as we have shown previously. Second, the dense coating of conductive polymers can be utilized to create entirely new conductive pathways between arbitrarily-defined neurons, microcircuits, or nervous system regions, thereby enabling researchers to effectively write new connections into living brains. Finally, this platform can be readily extended to a range of cell types, reactants (such as other redox-sensitive molecules), catalysts (such as enzymes or modulators), or reaction conditions (through modulating pH, light, chemical, redox, and electrical signals), which may enable investigators to establish high-content, specific, and seamless integration with living biological systems, beyond human design and assembly capabilities alone.

## Acknowledgements

National Science Foundation Future Manufacturing Program grant (award no. 2037164) supported method development for GTCA on neuronal membranes; the grant was awarded to K.D. and Z.B. based on the GTCA concept^8^. A.Z. gratefully acknowledges Professor Hollis Cline at Scripps Research and Dr. Shuo Han at Stanford University for helpful discussion, and Dr. Christina Newcomb at Stanford University for help with AFM imaging. Z.B. is a CZ Biohub investigator. K.Y.L. was supported by Stanford Bio-X. L.M. was supported by the Walter Benjamin Fellowship Program by the Deutsche Forschungsgemeinschaft (DFG 456522816). Part of this work was performed at Stanford University Cell Sciences Imaging Core Facility (RRID:SCR_017787). Part of this work was performed at the Stanford Nano Shared Facilities (SNSF), supported by the National Science Foundation under award ECCS-2026822.

## Author contributions

A.Z., K.D., and Z.B. conceived and designed the experiments. A.Z., K.Y.L., J.D., C.R. designed the molecular strategy. A.Z. developed the polymerization reactions, performed all sample preparation, imaging, SEM, UV-vis characterizations, and Raman spectroscopy. L.M. and A.Z. performed AFM imaging and analysis. C.S.K. and C.R. performed neuron culture. A.Z., K.D., and Z.B. analyzed all data, prepared figures and wrote the manuscript with edits from all authors. K.D. and Z.B. supervised all aspects of the work.

## Methods

### Plasmid constructs

The following constructs were designed in SnapGene 5.1.7 and cloned into AAV plasmids with the human Synapsin (hSyn) promoter. All sequences were confirmed with Sanger sequencing (Azenta).

HRP-CD2 or HRP(+): hSyn-IgK-FLAGx3-HRP-CD2-p2A-t2A-YFP

HRP-CD8: hSyn-IgK-FLAGx3-HRP-CD8-p2A-t2A-YFP

Apex2-CD2: hSyn-IgK-FLAGx3-Apex2-CD2-p2A-t2A-YFP

Apex2-CD8: hSyn-IgK-FLAGx3-Apex2-CD8-p2A-t2A-YFP

HRP(−): hSyn-IgK-FLAGx3-YFP-CD2-p2A-t2A-YFP

### Neuron culture, transfection, and antibody staining

Primary cultures of postnatal hippocampal rat neurons were prepared on 12-mm coverslips in 24-well plates as described previously^8^. Cells were transfected 6-7 days in vitro (DIV) with various constructs. For each coverslip to be transfected, a DNA-CaCl2 mix containing with the following reagents was prepared using calcium phosphate transfection kit (Invitrogen, 44-0052): 1 μg of plasmid DNA, 1 μg of salmon sperm DNA (Invitrogen, 15632-011), 1.875 μL 2M CaCl2, and sterile water added for a total volume of 15 μL. Finally, 15 μL of 2X HEPES-buffered saline (HBS) was added, and the resulting 30 μL mix was incubated at room temperature for 20 minutes. The growth medium from each well was removed, saved and replaced with 400 μL pre-warmed minimal essential medium (MEM, Invitrogen, 11095114), and the DNA-CaCl2-HBS mix was added dropwise into each well, and the plates were returned to the culture incubator for 60 minutes. Each coverslip was then washed three times with 1 mL of pre-warmed MEM, and placed back to the original neuronal growth medium. Neurons were used 4-6 days post-transfection.

To characterize membrane localization of HRP and Apex2, four days post-transfection, cultured neurons expressing different constructs were stained using two protocols, and imaged with Leica TCS SP8 confocal laser scanning microscope.

#### Non-detergent-permeabilized staining

Cultured neurons were washed 3 times with 1 mL of pre-warmed serum free Neurobasal medium (Gibco, 21103049) supplemented with 4% B-27 (Gibco, 17504044) and 2 mM Glutamax (Gibco, 35050061), fixed in 4% paraformaldehyde (PFA) at room temperature for 15 min, and washed 3 times with PBS. Cells were blocked with PBS containing 5% normal goat serum (Jackson ImmunoResearch, 005-000-121) at room temperature for 30 min, and then stained with primary antibody against FLAG (DDDDK tag, Abcam, ab1162) at 1:200 dilution with 5% goat serum at 37 °C for 1 h, and washed 3 times with PBS. Cells were then stained with Alexa 647 Goat anti-rabbit secondary antibody (Abcam, ab150087) at 1:500 dilution with 5% goat serum at 37 °C for 1 h, and washed 3 times with PBS. Cells were finally permeabilized in PBS containing 5% goat serum and 0.03% Triton X100 at room temperature for 10 min, and mounted on slides using VECTASHIELD hardSet antifade mounting medium with DAPI (Vector Laboratories).

#### Detergent-permeabilized staining

Cultured neurons were washed and fixed in 4% PFA as described above. Cells were then blocked and permeabilized with PBS containing 5% goat serum and 0.03% Triton X100 at room temperature for 1 h, and then stained with primary antibody and secondary antibody, and mounted with DAPI as described above.

### Evaluation of peroxidase activity

#### Amplex UltraRed staining

Live neurons in media were placed on ice for 3 min, and then incubated in 200 μL of 50 μM Amplex UltraRed (Invitrogen, A36006) in ice-cold Tyrode’s solution (125 mM NaCl, 2 mM KCl, 2 mM MgCl_2_, 2 mM CaCl_2_, 30 mM glucose, 25 mM HEPES; titrated to pH 7.35 with NaOH and adjusted osmolarity to 298) containing 0.05 mM H_2_O_2_ for 5 min, and washed with 1 mL fresh ice-cold Tyrode’s and imaged with Leica SP8 confocal microscope within 10 min.

#### 3,3’,5,5’-tetramethylbenzidine (TMB) staining

Cultured neurons were washed 3 times in media and fixed in 4% PFA at room temperature for 15 min, and washed 3 times with PBS. TMB solution was prepared by diluting the liquid TMB substrate containing 1.46 mM TMB, 2.21 mM H_2_O_2_ (Calbiochem, 3905420) 30 times in PBS. Fixed neurons were placed in the TMB solution for 10 min, and imaged with Leica SP8 confocal microscope within 10 min.

### Polymerization reactions

For polymerization reactions, cells were incubated in a mixture of polymer precursor solutions and hydrogen peroxide solution for 30 min and washed 3 times in Tyrode’s solution. All solutions were prepared fresh each time. 3 mM aniline dimer solution was prepared by dissolving 5.5 mg of N-phenyl-p-phenylenediamine (Sigma-Aldrich, 241393) in 10 mL of Tyrode’s solution ~20 h at room temperature, using a magnetic stir bar. 10 mM DAB solution was prepared by dissolving 36 mg of 3,3’-Diaminobenzidine tetrahydrochloride hydrate (Sigma-Aldrich, D5637) in 10 mL of Tyrode’s solution. The pH of the solution was adjusted to 7.35 by 1 M NaOH solution. When ready to perform the reactions, the above solutions were filtered with 0.45 μm syringe filters (Fisher Scientific), and H_2_O_2_ (EMD Millipore, 386790) was added to the solutions for a final concentration of 0.05 mM.

### Scanning electron microscopy (SEM)

Neurons were fixed with EM fix (2% glutaraldehyde (Electron Microscopy Sciences, 16020) with 4% PFA in 0.1 M Na Cacodylate buffer (Electron Microscopy Sciences, 11652)) at room temperature for 1 h, then washed 3 times in PBS. After reactions, cells were returned to EM fix and stored at 4 °C for 1 h, and then washed 3 times with 0.1 M Na Cacodylate buffer for 10 min each time. The buffer was then replaced with 1% aqueous Osmium Tetroxide (Electron Microscopy Sciences, 19192), at room temperature for 1 h, and washed with DI water twice for 10 min each time. Samples were then dehydrated in a series of ethanol washes (50%, 70%, 95%, 100%, 100%) for 10 min each, and then incubated in a series of hexamethyldisilazane (HMDS):ethanol mixtures (1:9, 2:8, 3:7, 4:6, 5:5, 6:4, 7:3, 8:2, 9:1) for 5 min each, followed by 100% HMDS twice for 20 min each time. The HMDS was then removed and the coverslips were dried overnight in a chemical hood. The samples were mounted with double-sided carbon tapes (Ted Pella, 16085-1) on aluminium tabs (Ted Pella, 16084-1), and coated with 4 nm gold with Leica EM ACE600 sputter coater, and imaged with a Zeiss Sigma SEM.

### Atomic force microscopy (AFM)

Morphology and modulus mappings were collected using a Bruker BioScope Resolve BioAFM with a SCANASYST-FLUID+ probe (Bruker) with nominal spring constant of 0.7 N/m and a tip radius of 2 nm on pre- and post-reacted, fixed neurons in Tyrode’s solution at room temperature. Nanomechanical maps were recorded in the Peak Force Quantitative Nanomechanics (PF-QNM) mode while scanning across the cell membrane for the height images. All force modulation measurements were performed at a setpoint of ~1.2 nN with a Peak Force frequency of 0.5 kHz and amplitude of 300 nm. The scan resolution was set to 256×256 pixel with a scan-rate of 0.5 Hz. The data was evaluated and depicted with Gwyddion SPM software. The histograms of the modulus channel were generated via a 1D statistical function embedded in Gwyddion.

### Ultraviolet-visible (UV-Vis) spectrophotometry and Raman spectroscopy

Post-reacted, fixed neurons on glass coverslips were washed 3 times in PBS and 3 times in DI water, air-dried and mounted onto the sample holders. UV-Vis spectrophotometry was performed with Agilent Cary 6000i spectrophotometer. Raman spectroscopy was performed with Horiba XploRA Confocal Raman Microscope with a 785 nm laser using a 100x Olympus MPLan N objective.

### Cell viability assay

To examine cell viability of neurons post-reaction, coverslips with live HRP(+) neurons were fixed on the sample platform of a Leica DMi8 microscope and imaged with the YFP signal. After polymerization (or fixation in 4% PFA at room temperature for 15 min and permeabilization with 0.03% Triton X100 at room temperature for 15 min as the positive control), dead cells were stained red with 4 μM ethidium homodimer (EthD-1, Invitrogen, L3224) in Tyrode’s solution at room temperature for 30 min, and imaged in the red channel.

### Data analysis

ImageJ software was used for imaging analysis, including fluorescent intensity and brightness measurements. For cell viability tests, the yellow and red channel images were concatenated, and the locations of cells expressing YFP were marked with white dots in the red channel. Spectroscopic results were plotted using Matlab R2022b. Statistical analyses for all data were performed with two-tailed unpaired t-test using GraphPad Prism 9. Chemical structures were prepared using ChemDraw 21.0.

