## Supplementary Information for "In situ genetically targeted chemical assembly of polymers on living neuronal membranes"

**In situ genetically targeted chemical assembly of polymers on living neuron membrane**

This file contains:

Supplementary Figures S1 – S8

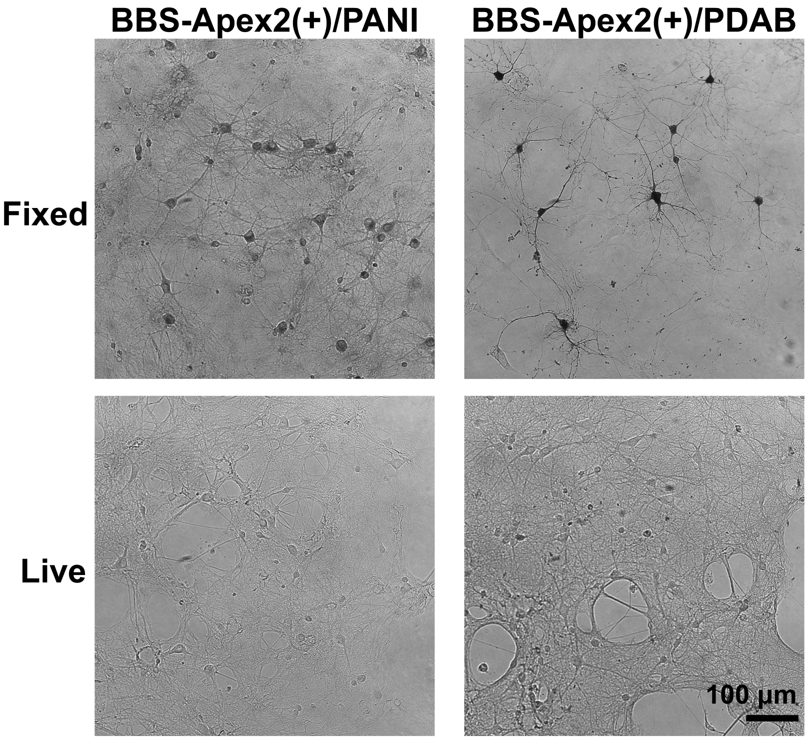


**Supplementary Figure S1** Comparison of polymerization on fixed and live neurons transfected with AAVdj viruses encoding hSyn-BBS-Apex2-YFP using reaction conditions as described in Liu et al., Science 367, 1372-1376 (2020). The assembled polymers form dark aggregates in cells. In fixed neurons, wherein cell membranes were permeabilized during polymerization, both intracellular and extracellular Apex2 catalyzed polymerization. In live neurons, wherein cell membranes remained intact during polymerization, extracellular Apex2 more robustly catalyzed polymerization; fixed BBS-Apex2 neurons were darker than live neurons, consistent with the interpretation that Apex2 in previous work was expressed intracellularly to a substantial extent. Scale bar: 100 µm.

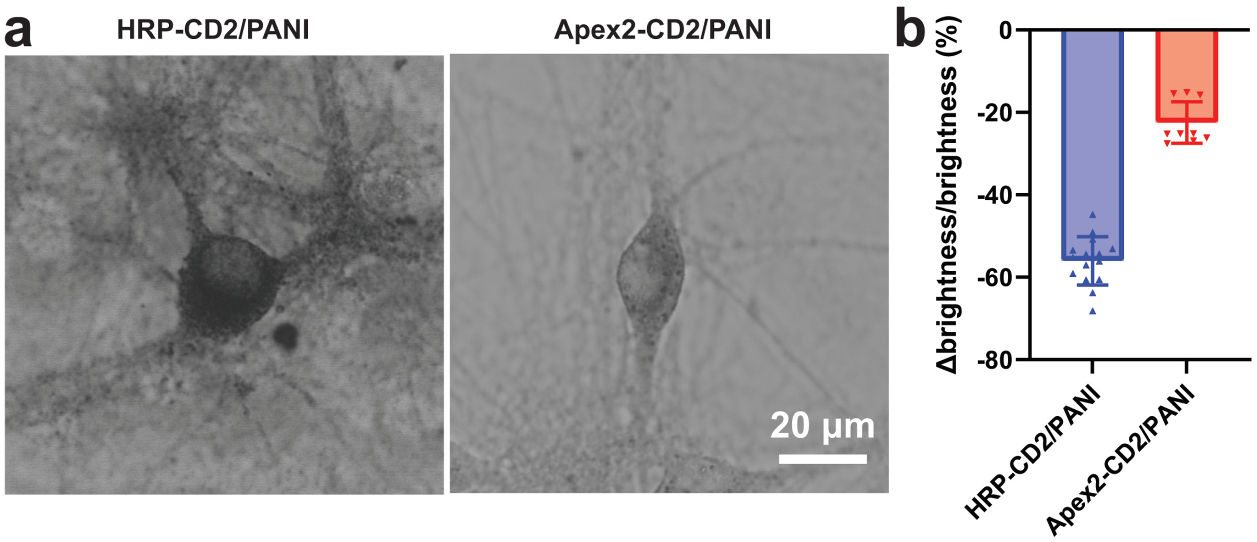


**Supplementary Figure S2** Comparison of PANI polymerization reactions on cells expressing HRP-CD2 and Apex2-CD2. (**a**) Bright field images of live neurons with and without PANI deposition. (**b**) Statistical comparison of the ratio (expressed as %) of brightness difference between neuron and background ("Δbrightness"), compared to background brightness. N = 15 cells for HRP-CD2/PANI, N = 10 cells for Apex2-CD2/PANI. Values are means ± s.d.. Polymerization rate on HRP-CD2 cells are faster than on Apex2-CD2 cells.

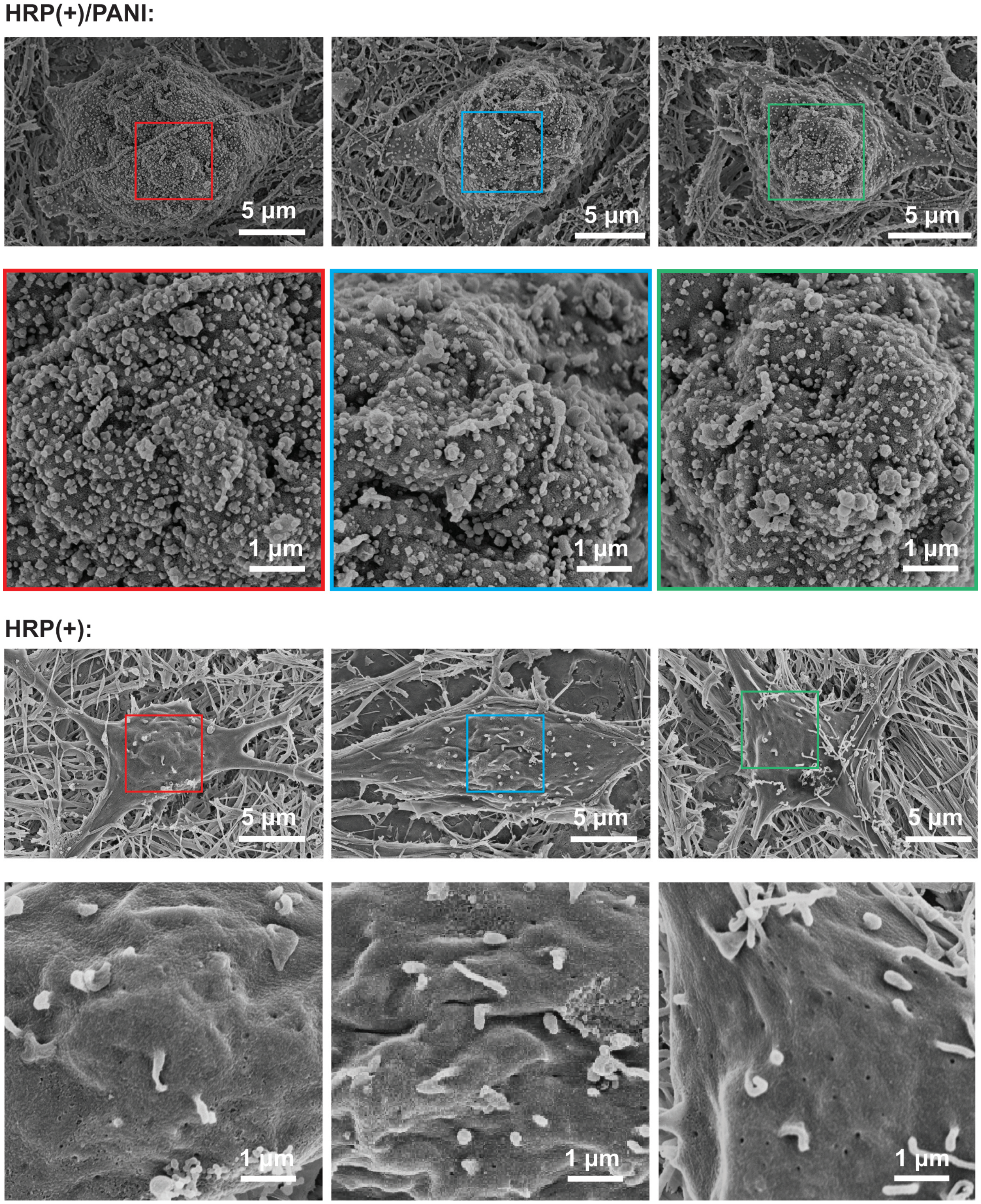


**Supplementary Figure S3** Additional SEM images of HRP(+)/PANI and HRP(+) neurons.

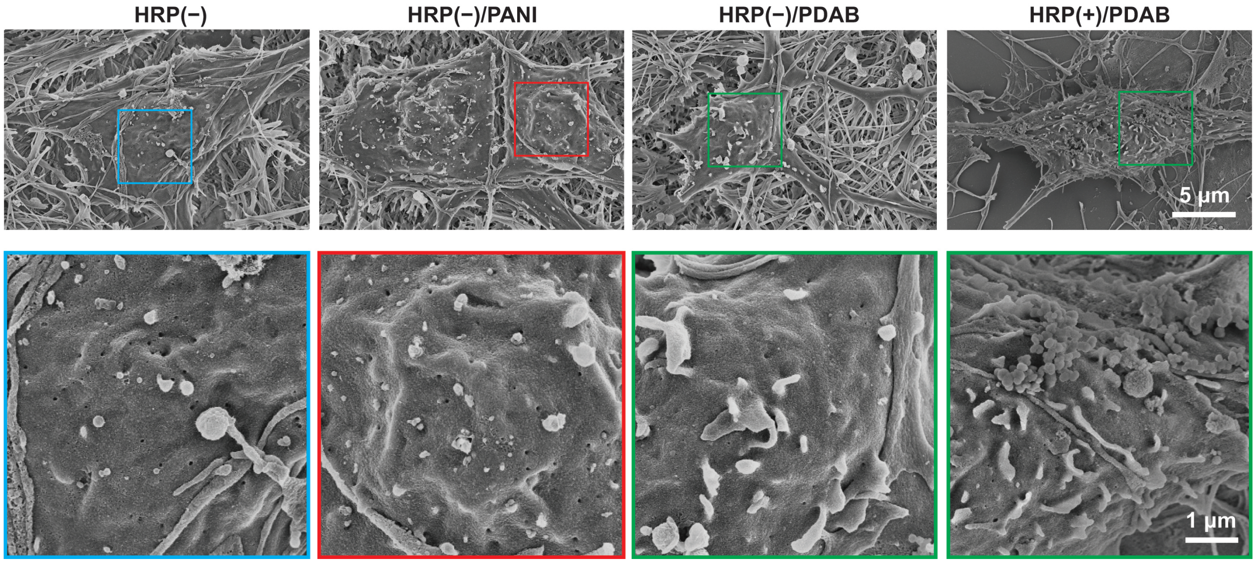


**Supplementary Figure S4** SEM images of HRP(−) neurons with and without polymerization reactions, and HRP(+) neurons with PDAB.

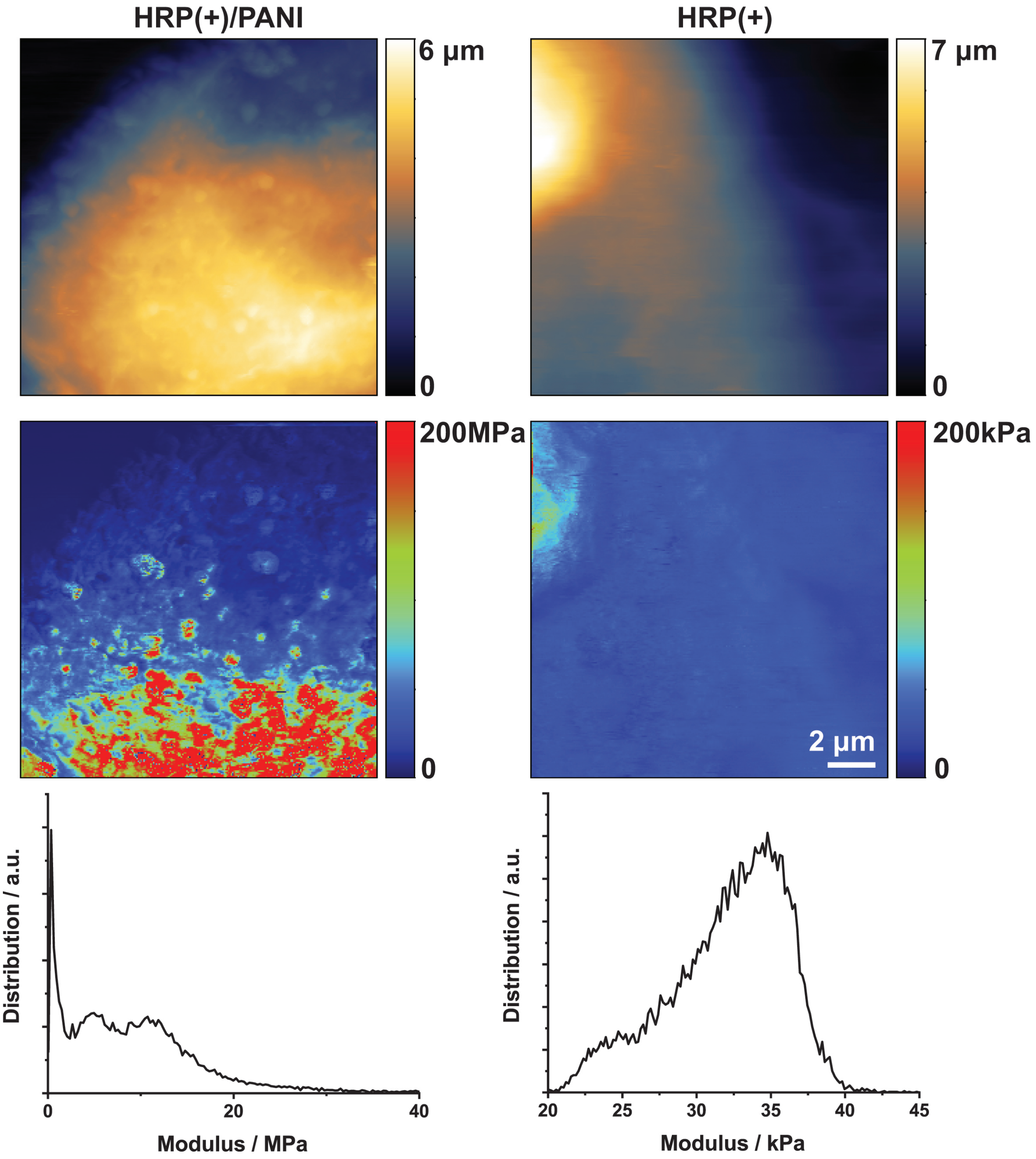


**Supplementary Figure S5** Top, original AFM height and modulus images of the panels in Fig. 3d. Bottom, distribution of the modulus in the original images.

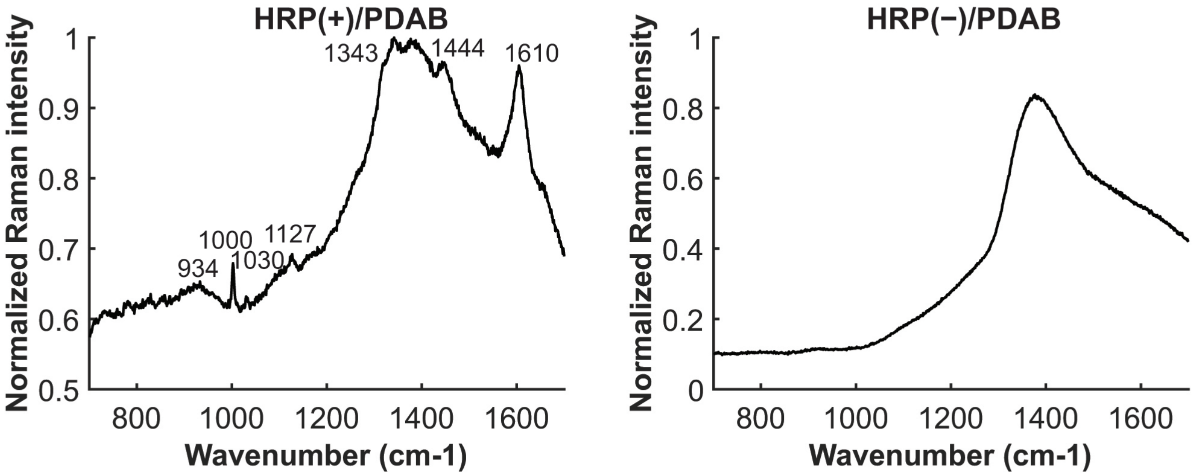


**Supplementary Figure S6** Raman spectroscopy analysis of the HRP(+) and HRP(−) after PDAB deposition.

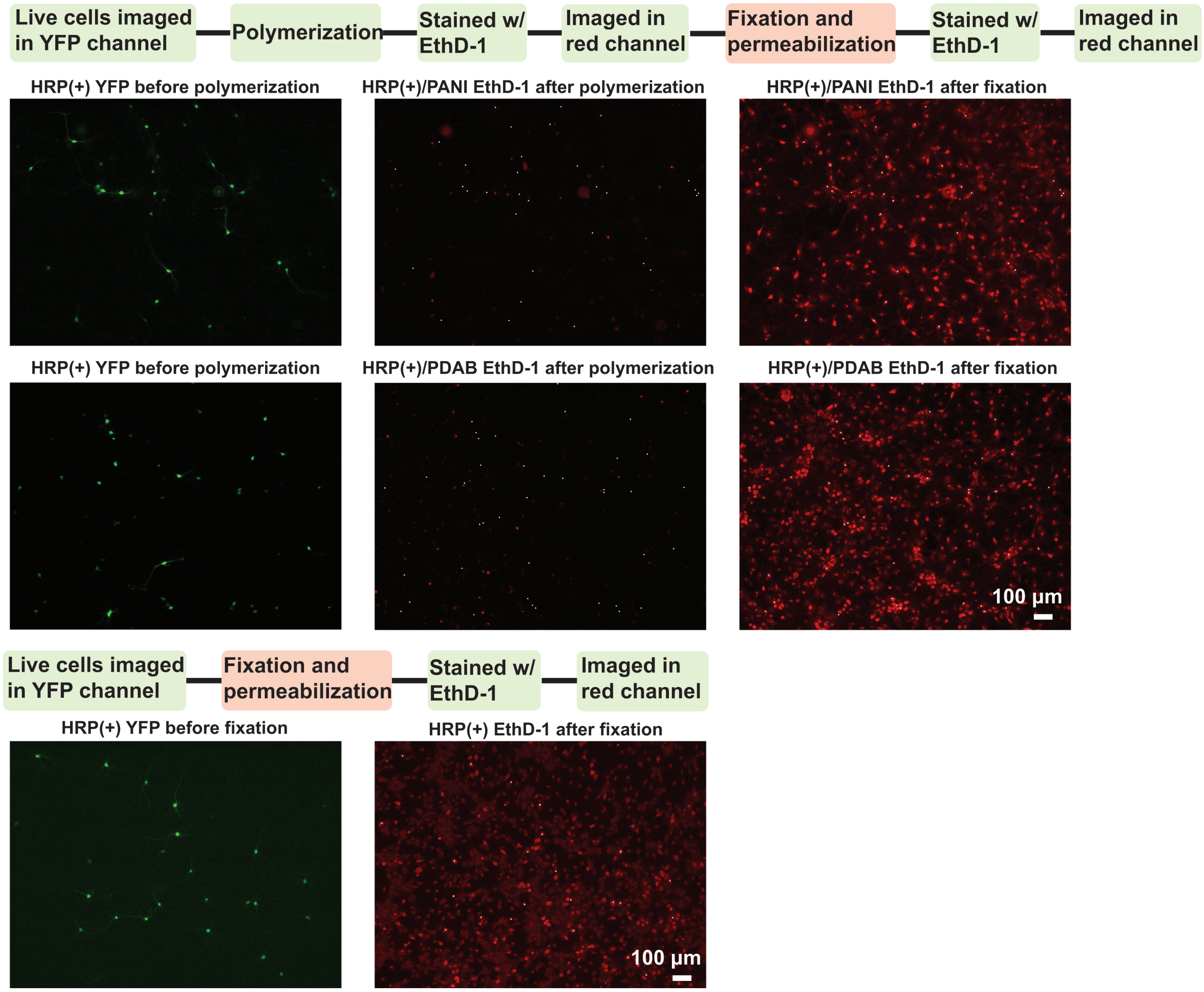


**Supplementary Figure S7** Positive control for the cell viability tests. Top, live HRP(+) cells were imaged with YFP, and after PANI or PDAB deposition, were stained and imaged with ethidium homodimer (EthD-1), a red fluorophore that only stains dead cells with damaged membrane. The cells were then fixed and stained with EthD-1 again. The white dots in the EthD-1 images mark the locations of the HRP(+) cells. No white dot overlays with the red cells in the images in the middle, indicating HRP(+) neurons remain viable after polymerization. All white dots overlay with the red cells in the images on the right, indicating that EthD-1 properly labeled all dead cells. Bottom, live HRP(+) cells were imaged with YFP (left), and after fixation and permeabilization, were stained with EthD-1 (right). All white dots in the EthD-1 image overlay with the red cells, indicating that EthD-1 properly labeled all dead cells.

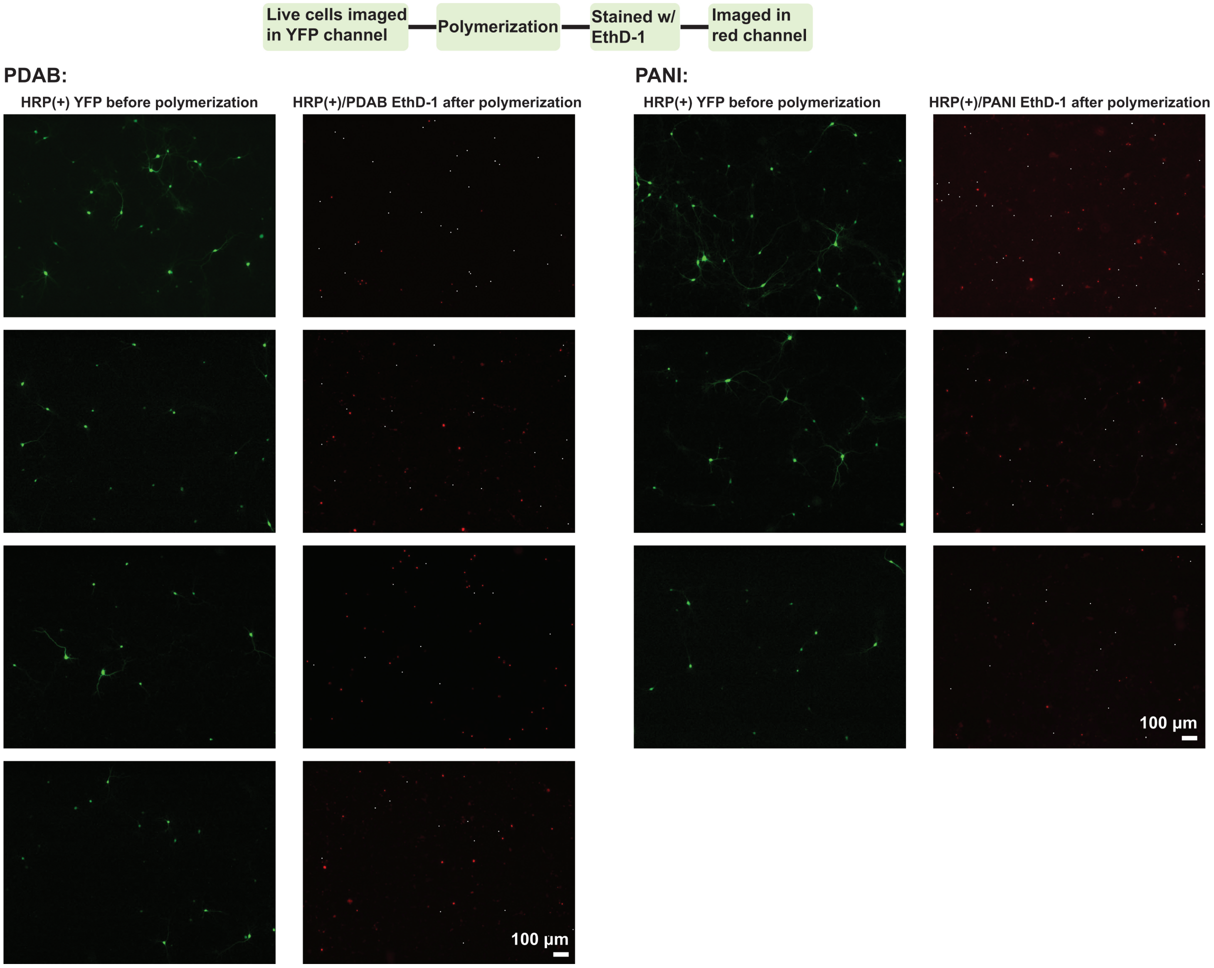


**Supplementary Figure S8** Additional cell viability tests of HRP(+) cells after PDAB and PANI deposition. N = 5 coverslips were used for each polymerization condition. One field of view was imaged on each coverslip. All HRP(+) neurons remain viable after polymerization.
